# PhyloCSF++: A fast and user-friendly implementation of PhyloCSF with annotation tools

**DOI:** 10.1101/2021.03.10.434297

**Authors:** Christopher Pockrandt, Martin Steinegger, Steven L. Salzberg

## Abstract

**Summary:** PhyloCSF++ is an efficient and parallelized C++ implementation of the popular PhyloCSF method to distinguish protein-coding and non-coding regions in a genome based on multiple sequence alignments. It can score alignments or produce browser tracks for entire genomes in the wig file format. Additionally, PhyloCSF++ annotates coding sequences in GFF/GTF files using precomputed tracks or computes and scores multiple sequence alignments on the fly with MMseqs2.

**Availability:** PhyloCSF++ is released under the AGPLv3 license. Binaries and source code are available at https://github.com/cpockrandt/PhyloCSFpp. The software can be installed through bioconda. A variety of tracks can be accessed through ftp://ftp.ccb.jhu.edu/pub/software/phylocsfpp/.

## 1 Introduction

Two decades after the first human genome was sequenced, previously unknown genes in human are still being found [7]. For many less-studied organisms the situation is even more uncertain, as their protein-coding gene catalogues remain sparse.

An established method that uses evolutionary conservation to detect protein-coding regions is PhyloCSF [4]. It takes a multiple sequence alignment (MSA) and a phylogenetic tree of related species as input and computes a score, based on codon substitution rates and codon frequencies, reflecting the likelihood for the region to be coding or non-coding. As examples of its recent application, PhyloCSF scores (available as genomic tracks) were used to identify 133 novel protein coding genes in human [5] and to investigate the conservation of the newly emerged ORF10 in *SARS-CoV2* [2]. Currently, there are tracks available for *H.sapiens, M.musculus, G.gallus, C.elegans, D.melanogaster, A.gambiae* and *SARS-CoV-2.*

Here we present the free open-source software PhyloCSF++, a fast multi-threaded implementation of PhyloCSF that additionally to scoring alignments directly produces genome tracks, and allows users to annotate GFF files with PhyloCSF scores. We used PhyloCSF++ to generate tracks for 5 additional species and made them available for the UCSC genome browser [3]^1^.

## 2 Methods

The original PhyloCSF tool was developed in OCaml and requires older versions of the OCaml compiler and libraries, making it challenging to compile it from source. Since 2019 it became installable through bioconda lowering the barrier for users to run it. However, no stand alone binary version of it is available. Computing tracks and posterior probabilities requires additional scripts [5] and coding, and multi-threading is not supported.

PhyloCSF is a valuable method that after 10 years still plays an important role in the discovery of novel genes, hence we created PhyloCSF++, a new implementation of the algorithm in C++ that is more efficient and that adds parallelization. Multi-threading allows one to process large MSAs of eukaryotic genomes. It can be installed on any UNIX system with *gcc* ≥ 4.9 and *cmake,* and only depends on the GNU scientific library [1]. Statically linked binaries are available for download, or can be installed via bioconda.

## 3 Results

### Score computation

PhyloCSF++ offers all three PhyloCSF modes (fixed, mle and omega) that compute the likelihood of a region to be coding or noncoding. All three modes are faster than PhyloCSF and fully parallelized. In a benchmark on a subset of alignments from the 100-way multiz alignment of GRCh38 using the 58mammals model, PhyloCSF++ was faster by a factor of 1.3, 2.2 and 2.4 for all three modes with a single thread compared to PhyloCSF (see Fig. 1d). The maximum likelihood estimation as well as the omega mode (that only requires a phylogenetic tree and no training data) have randomized algorithms and can lead to minor differences in computed scores compared to the reference implementation.

**Fig. 1.**
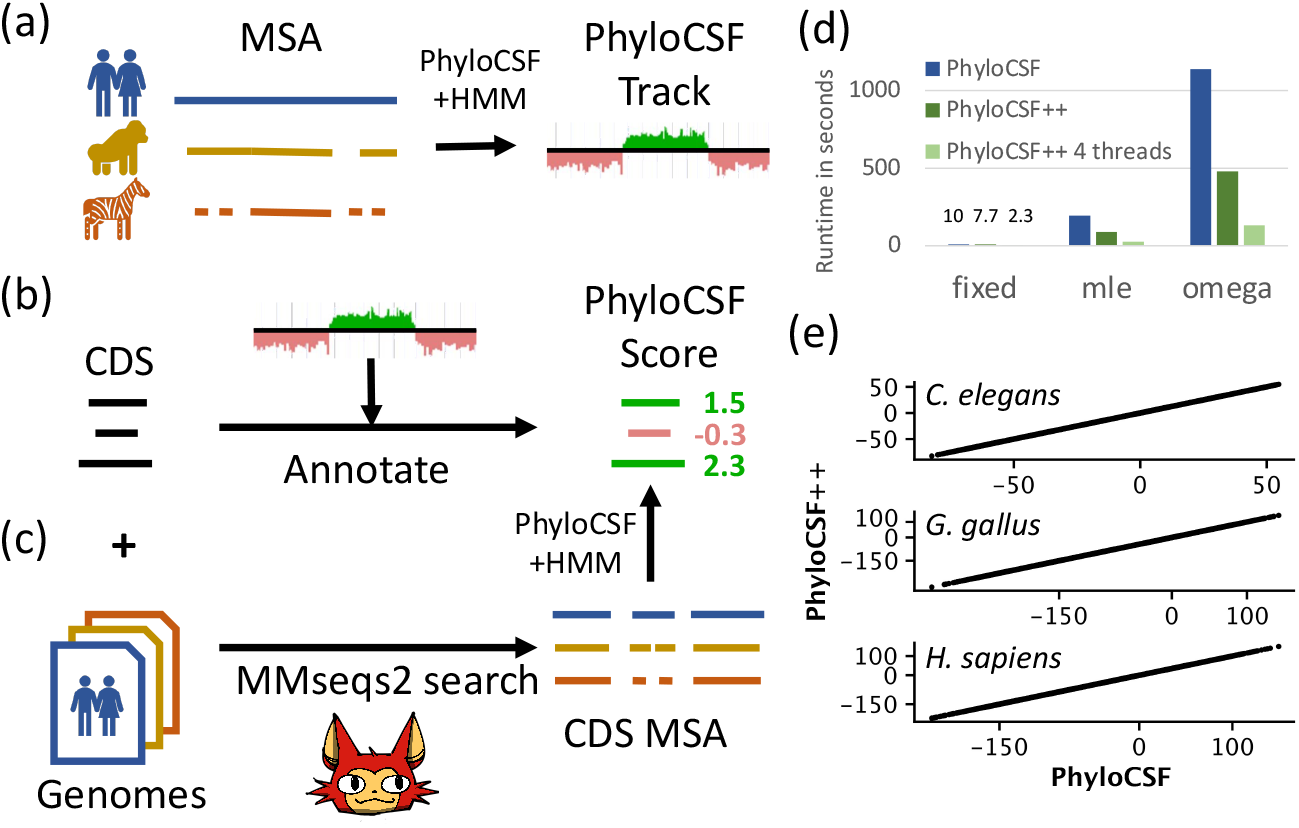
PhyloCSF++. (a) The command *build-tracks* of PhyloCSF++ takes a whole-genome MSA of related species as input and produces six tracks, one for each reading frame. Additionally, a confidence track is outputted. Each track is stored as wig file and consists of normalized posterior probability scores within the range ±15 for each codon of the input MSA. Positive scores indicate coding and negative scores indicate non-coding regions. (b) The command *annotate-with-tracks* scores coding sequences (CDS) from a GFF file using PhyloCSF tracks. (c) Alternatively, CDS can be scored without tracks using MMSeqs2 [8] and a set of genome sequences. MMSeqs2 will perform a search and generate a MSA of all CDS, which then will be scored by PhyloCSF++. (d) Speed comparison for generating tracks of PhyloCSF (blue) and PhyloCSF++ single threaded (dark green) and multi threaded (green) for the first 50 alignment blocks with a minimum sequence length of 50 of chr22 of the 100way alignment of GRCh38 with the 58mammals model. (e) Comparison of raw PhyloCSF and PhyloCSF++ tracks of ce11, galGal6 and hg38 (up to down). Both produce nearly equal scores with a Pearson correlation coefficient of 1.0.

### Track building

In contrast to the original implementation of PhyloCSF, PhyloCSF++ offers a direct way to build tracks. For C.elegans, G.gallus and H.sapiens we recomputed the tracks and got effectively the same scores for all codons and all frames, with a Pearson correlation coefficient of 1.0 for raw scores and 0.999 for smoothed scores. We used the same models (8worms, 53birds, 58mammals) and alignments as the original PhyloCSF tracks [5] (see PhyloCSF Track Hub). Fig. 1e shows the dot plot of 10^7^ sampled positions across all frames for each species.

### GFF annotation with tracks

To simplify the evaluation of PhyloCSF scores, the tracks can be used to annotate CDS features in a GFF file. For each CDS feature and its corresponding transcript, PhyloCSF scores as well as confidence scores are computed.

### GFF annotation with MMseqs2

If tracks or a whole-genome MSA are not available, a GFF file can be annotated by aligning only the CDS to a set of reference genomes on the fly using MMseqs2. MMseqs2 computes and chains MSAs directly for the transcripts of interest without the need for a whole-genome multiple sequence alignment, which would require a much greater computing resource than aligning only the CDS.

### Tracks

We generated new tracks for the following species: rat (rn6), fugu (fr3), stickleback (gasAcu1), tarsier (tarSyr2) and yeast (sacCer3). Each of these tracks is available for download and also accessible through the UCSC genome browser.

## Conclusion

PhyloCSF++ was developed to make the discovery of novel genes for many species faster and easier. Additional features to the original PhyloCSF software allow users to directly build tracks from whole-genome MSAs. Furthermore, PhyloCSF++ can directly annotate GFF files using computed tracks, or if no whole-genome MSAs are available, quickly compute MSAs using MMseqs2.

In summary, we hope that the software will prove helpful to detect novel protein sequences in already well-studied genomes and also prove useful for the large and growing number of new genomes [6].

## Footnotes

1 ftp://ftp.ccb.jhu.edu/pub/software/phylocsfpp/hub.txt

